# Aberrant epigenetic and transcriptional events associated with breast cancer risk

**DOI:** 10.1101/2021.09.14.460320

**Authors:** Natascia Marino, Rana German, Ram Podicheti, Douglas B. Rush, Pam Rockey, Jie Huang, George E. Sandusky, Constance J. Temm, Sandra K. Althouse, Kenneth P. Nephew, Harikrishna Nakshatri, Jun Liu, Ashley Vode, Sha Cao, Anna Maria Storniolo

**Affiliations:** Susan G. Komen Tissue Bank at the IU Simon Comprehensive Cancer Center, Indianapolis, IN 46202, USA; Department of Medicine, Indiana University School of Medicine, Indianapolis, IN 46202, USA; Center for Genomics and Bioinformatics, Indiana University, Bloomington, IN 47405, USA; Pathology and Laboratory Medicine, Indiana University School of Medicine, Indianapolis, IN 46202, USA; Department of Biostatistics and Health Data Science, Indiana University School of Medicine, Indianapolis, IN 46202, USA; Department of Anatomy, Cell Biology, & Physiology, Indiana University, Bloomington, IN 47405, USA; Department of Surgery, Indiana University School of Medicine, Indianapolis, IN 46202, USA

**Author notes:** Corresponding author: Natascia Marino, PhD; Department of Medicine, Hematology/Oncology Division; Indiana University School of Medicine; 980 W. Walnut St, R3-C238, Indianapolis, IN 46202.

**Keywords:** Cancer risk, transcriptome, DNA methylation, normal breast

## Abstract

**Background:** Genome-wide association studies have identified several breast cancer susceptibility loci. However, biomarkers for risk assessment are still missing. Here, we investigated cancer-related molecular changes detected in tissues from women at high risk for breast cancer prior to disease manifestation. Disease-free breast tissue cores donated by healthy women (N=146, median age=39 years) were processed for both methylome (MethylCap) and transcriptome (Illumina’s HiSeq4000) sequencing. Analysis of tissue microarray and primary breast epithelial cells was used to confirm gene expression dysregulation.

**Results:** Transcriptomic analysis identified 69 differentially expressed genes between women at either high and those at average risk of breast cancer (Tyrer-Cuzick model) at FDR<0.05 and fold change≥2. The majority of the identified genes were involved in DNA damage checkpoint, cell cycle, and cell adhesion. Two genes, FAM83A and NEK2, were overexpressed in tissue sections (FDR<0.01) and primary epithelial cells (*p*<0.05) from high-risk breasts. Moreover, 1698 DNA methylation aberrations were identified in high-risk breast tissues (FDR<0.05), partially overlapped with cancer-related signatures and correlated with transcriptional changes (*p*<0.05, *r*≤0.5). Finally, among the participants, 35 women donated breast biopsies at two time points, and age-related molecular alterations enhanced in high-risk subjects were identified.

**Conclusions:** Normal breast tissue from women at high risk of breast cancer bears molecular aberrations that may contribute to breast cancer susceptibility. This study is the first molecular characterization of the true normal breast tissues and provides an opportunity to investigate molecular markers of breast cancer risk, which may lead to new preventive approaches.

## BACKGROUND

Genetic and epigenetic alterations in breast cancer (BC) have been widely investigated. However, when, during the carinogenesis process, these events first emergeremains unknown. The identification of molecular aberrations associated with BC development can provide a conceptual framework for a deeper understanding of this complex disease.

Genome-wide association studies (GWAS) have detected more than 170 genomic loci harboring common variants associated with BC risk including modifier alleles with high (e.g., BRCA1, BRCA2, TP53, PTEN) to moderate penetrance (e.g., BRIP1, CHEK2, ATM, and PALB2) (1-4). Nevertheless, many variants are located in noncoding or intergenic regions and their functional role in cancer transformation remains largely unknown. Recently, transcriptome-wide association studies were used to integrate GWAS and gene expression datasets and identified 154 genes whose predicted expression associated with the risk for BC (5-9). However, these studies drew data from the Genotype-Tissue Expression (GTEx) project, where the use of autopsy-derived normal breast tissues may make the breast-specific transcriptome profilings questionable. The relative lack of molecular profiling of normal breast tissue from subjects who are disease-free makes the field challenging.

Many studies searching for cancer biomarkers have identified gene expression signatures, epigenetic signatures, loss of heterozygosity and allelic imbalance resulting from the development of malignancy (10). Among the molecular processes linked with cancer, DNA methylation has a keyrole in early cancer development through a process known as epigenetic reprogramming (11), potentially leading to silencing and loss of expression of tumor suppressor genes (12), and genomic instability (13).

Here, we performed an integrated analysis of DNA methylation and transcriptome profiling of cancer-free breast tissues donated by healthy women at either average or high risk for BC. In addition to early epigenetic events, we identified two molecules overexpressed in high-risk breasts independently from DNA methylation changes and, therefore, potential markers of BC susceptibility. Moreover, using a subcohort of repeated breast tissues donation by the same donors, we confirmed that the molecular changes identified in high-risk subjects are age-independent. These findings will lead to a deeper understanding of BC susceptibility and also provide the scientific community with the molecular profiling of a true normal breast tissue.

## RESULTS

### Study cohort used to investigate molecular aberrations in association with breast cancer (BC) risk

To identify transcriptomic and epigenetic differences linked with BC risk, we analyzed cancer-free breast tissue cores donated by 146 healthy women (median age: 39 years) including 112 Caucasian, 24 African American and 10 Asian subjects (**Additional File 1: Table S1**). Out of 146 participants, 117 were pre- and 29 postmenopausal women. Tyrer-Cuzick model was employed to estimate the lifetime risk of developing BC and allocated the subjects to either high- (score≥20%, N=68) or average-risk group (score<20%, N=78) (**Fig. 1A, Table 1** and **Additional File 1: Table S1**).

**Figure 1:**
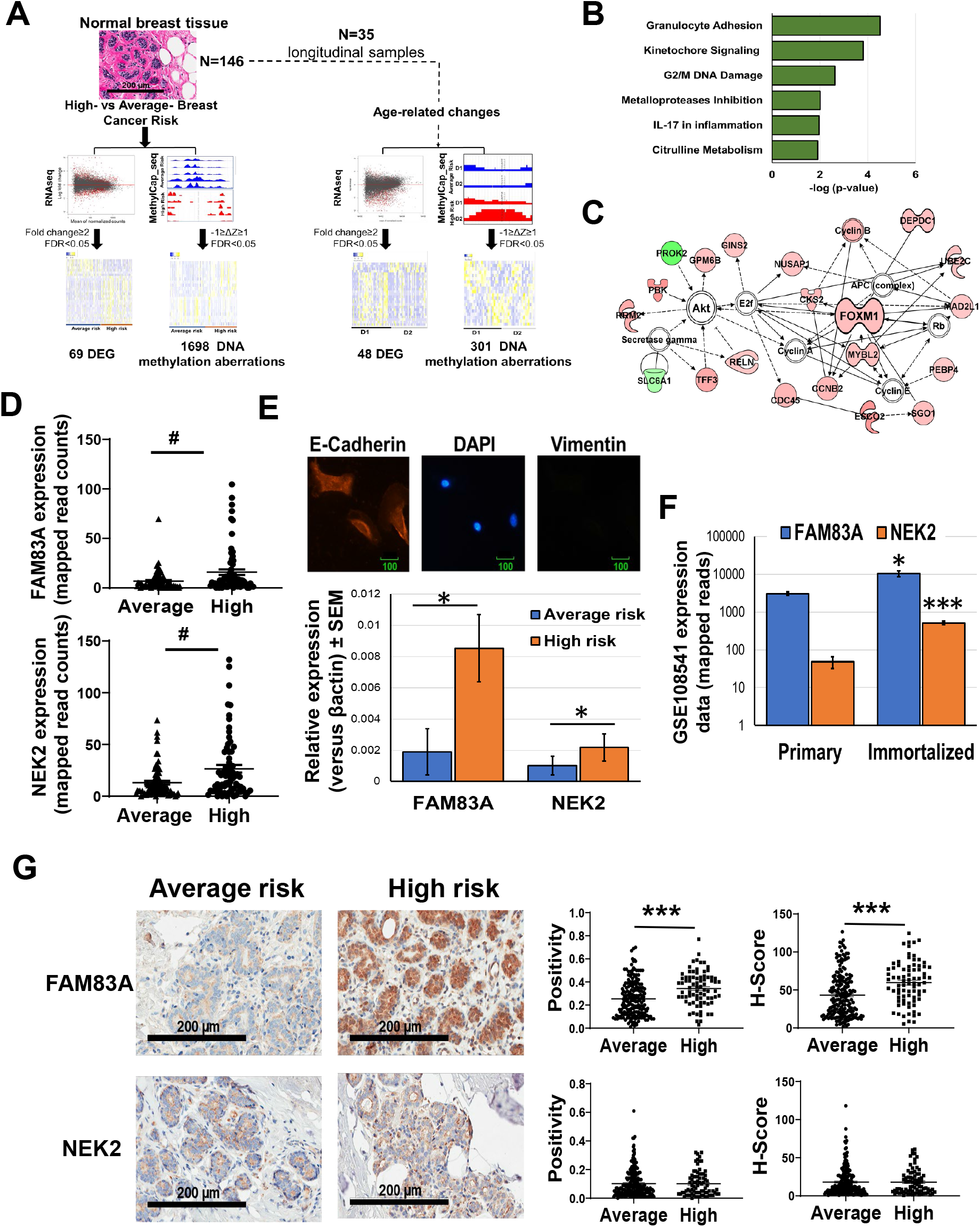
Transcriptome profiling of breast tissues from women at either high- or average risk of breast cancer. **A)** Schematics of the study design. Cancer-free breast tissue cores (N=146) were divided in either high-risk or average-risk group according to the Tyrer-Cuzick breast cancer risk evaluation score (20% used as threashold). The tissues were processed and analyzed for whole transcriptome and methylome profiling and differentially expressed genes (DEG) and differentially methylated sites between high- and average-risk samples were identified. Thirty five women (10 high risk and 25 average risk) donated also a second biopsy (D2) allowing to detect age-dependent aberrations. **B)** Pathway analysis of the transcripts differentially expressed (FDR<0.05) between average and high-risk breasts. **C)** Major molecular network of the differentially expressed transcripts between the two experimental groups. Genes upregulated in high-risk breasts are in red, while downregulated genes are in green. **D)** FAM83A and NEK2 transcription level in breast tissues from women at either average- or high-risk of developing breast cancer. **E**) Upper panel: Representative image of the immunofluorescence staining of primary breast epithelial cells with the epithelial marker, E-Cadherin (red), mesenchymal marker, Vimentin (green) as control, and nuclear staining, DAPI (blue). E-Cadherin and Vimentin staining of primary cells revealed that isolated primary cells are epithelial in nature. Lower panel: FAM83A and NEK2 expression in primary epithelial cells isolated from either average-risk (n=4) and high-risk breast (n=3). **F**) FAM83A and NEK2 expression in primary and h-TERT immortalized isogenic breast epithelial cells (n=7) from the GSE108541 dataset. **G**) Representative images of immunohistochemical staining for FAM83A and NEK2 are shown at 20X magnification. Staining quantification is expressed as positivity and H-score. Data are shown as mean ± standard error. #: FDR<0.005, *: *p*<0.05,\*\**p*<0.001, \*\*\**p*<0.0001. *P*value is calculated using either unpaired nonparametric Mann-Whitney test or nonparametric Wilcoxon test.

**Table 1:**
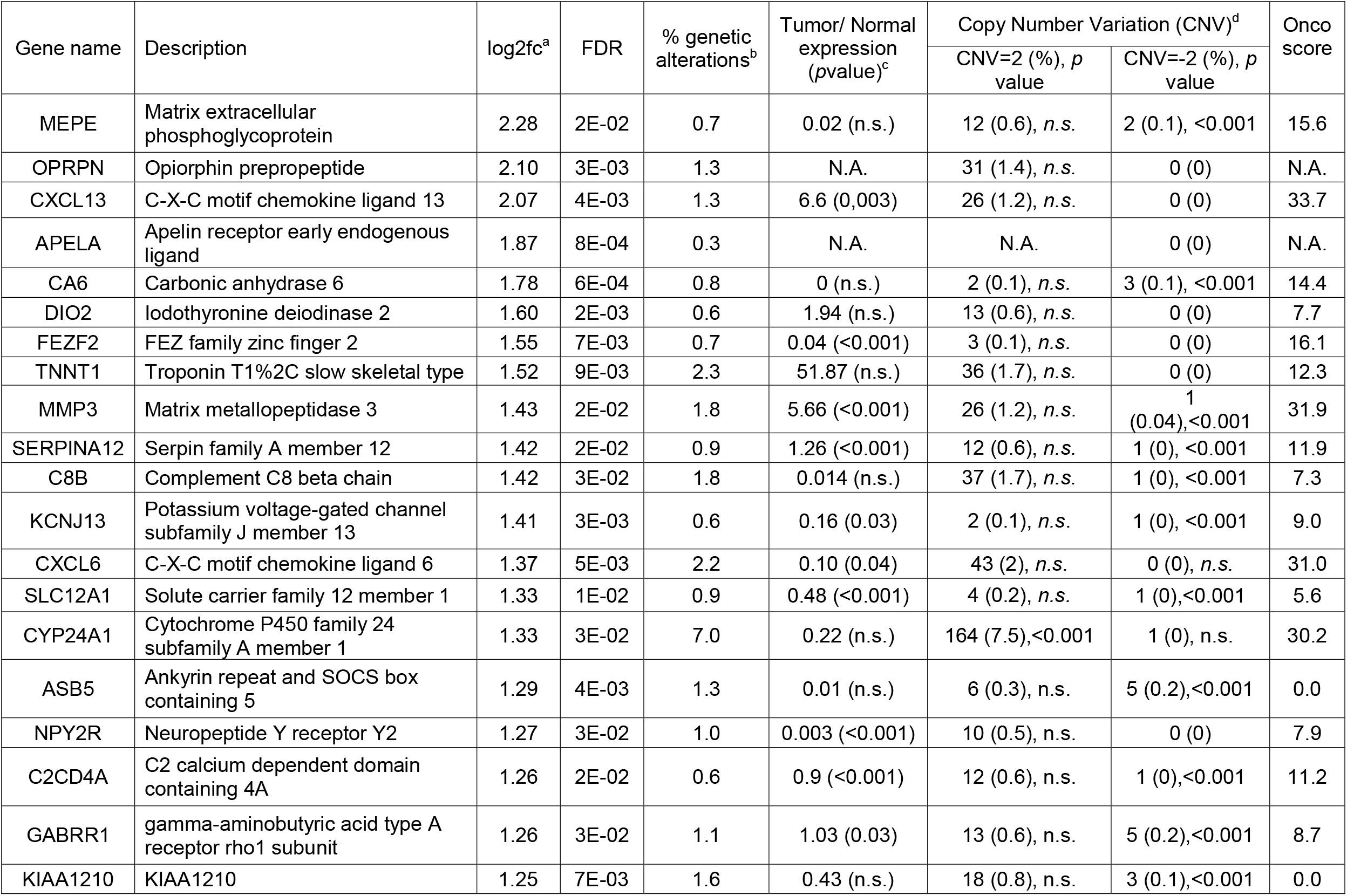
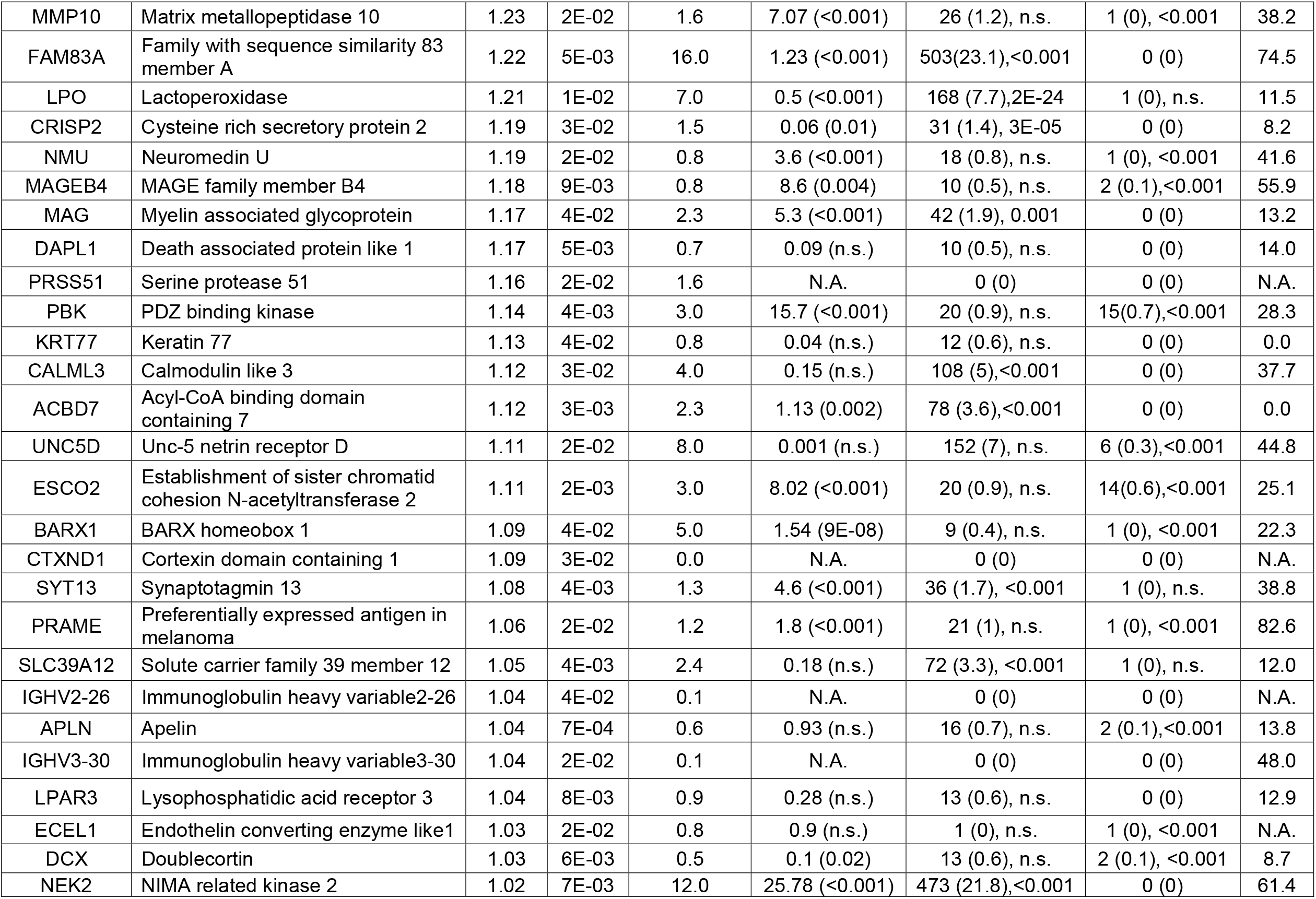
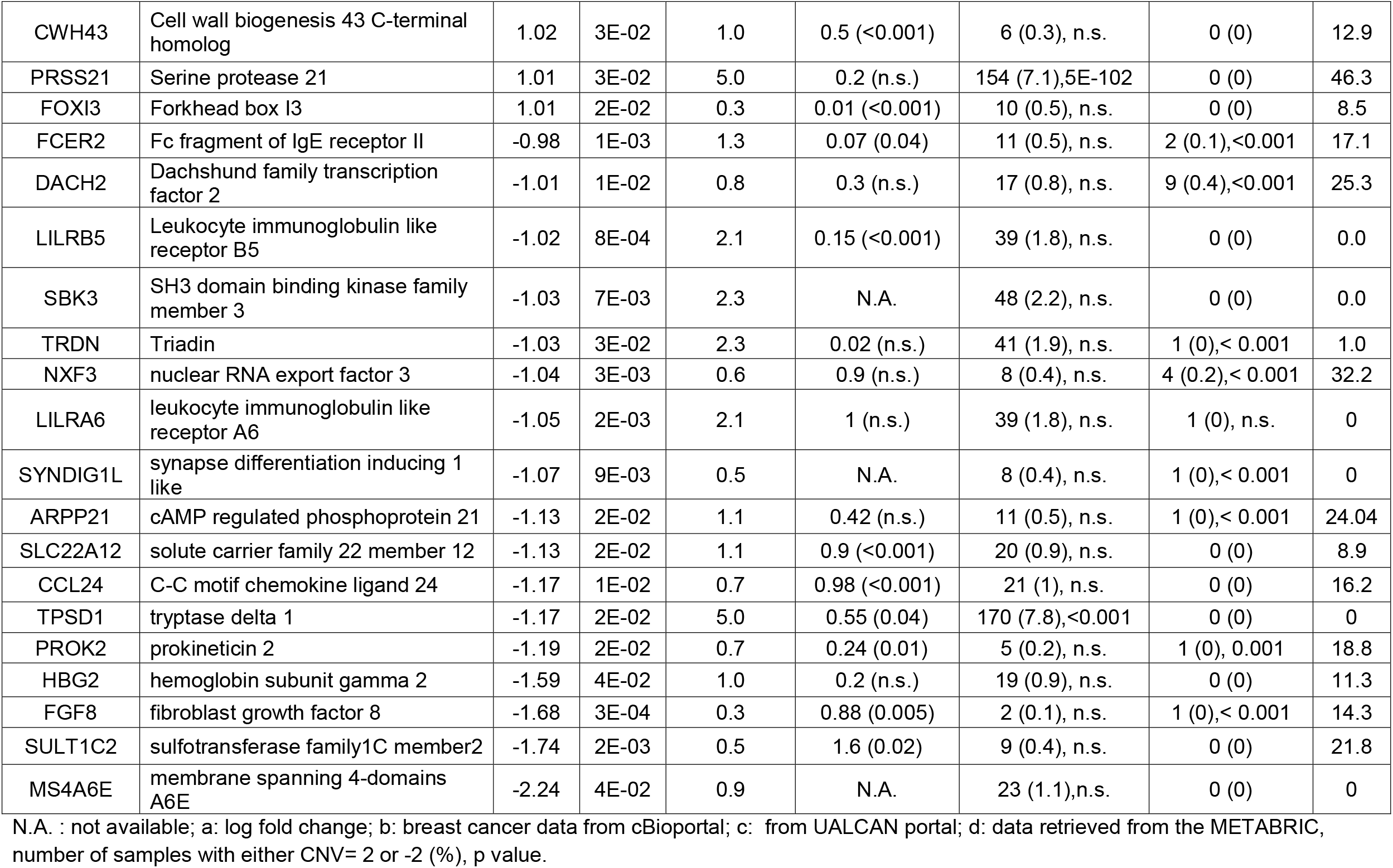
Gene expression differences in high- versus average-risk breasts (FC>2; FDR<0.05)

### Characterization of the transcriptome alterations in high-risk breast

We performed a transcriptome analysis of the fresh frozen disease-free breast tissue donated by all the participants. Differential expression analysis was performed using DESeq2. From a total of 22,344 genes, the differential expression analysis between high- and average-risk breasts revealed 1,874 transcripts to be significant at 5% false discovery rate (FDR). Of these, 1,798 transcripts also passed the cutoff of *t*-test *p*-value ≤ 0.05 (**Additional File 1: Table S2**). Sixty-nine genes, including 51 upregulated and 18 downregulated genes, were identified with a fold change ≥ 2 (**Table 1**). Because both groups consisted of non-malignant breast tissue, a limited number of differentially expressed genes was expected (14). Canonical pathway analysis revealed enrichment in pathways related to kinetochore signaling (*p*=1.3E-05), DNA damage checkpoint (*p*=0.0005), granulocytes adhesion (*p*=0.002), and the IL17 pathway (*p*=0.004) (**Figure 1B, Additional File 1: Table S3)**. Our data further confirm the previously described impact of dysregulated DNA damage in breast carcinogenesis (15). Molecular network analysis showed an enrichment in functional categories involved in cell cycle, DNA replication and repair (**Figure 1C, Additional File 1: Table S3**). One of the major molecular networks regulating cell cycle is centered around AKT and the transcription factor FOXM1 (16).

Except for DCX, the transcriptional changes detected between high- and average-risk breasts listed in Table 2 are independent of both racial background and menopausal status of the tissue donors (**Additional File 1: Table S4** and **Additional File 2: Fig. S1**).

**Table 2:** The 20 most differentially methylated regions between the high- and average-risk breast tissues.

| Genomic Locus | Overlapping Gene Feature | Gene Name | Description | $\Delta Z^{\#}$ | FDR |
| --- | --- | --- | --- | --- | --- |
| Chr8:120,669,501 | Intron | SNTB1 | syntrophin beta 1 | 2.4 | 7E-53 |
| Chr18:6,803,751 | Intron | ARHGAP28 | Rho GTPase activating protein 28 | 2.0 | 3E-35 |
| Chr6:12,944,751 | Intron | PHACTR1 | phosphatase and actin regulator 1 | 1.9 | 1E-31 |
| Chr21:30,743,501 | promoter | KRTAP21-4P | keratin associated protein 21-4 2C pseudogene | 1.9 | 6E-31 |
| Chr2:115,663,751 | Intron | DPP10 | dipeptidyl peptidase like 10 | 1.8 | 2E-30 |
| Chr4:87,111,001 | Intron | AFF1 | AF4/FMR2 family member 1 | 1.8 | 2E-29 |
| Chr3:33,638,251 | Intron | CLASP2 | cytoplasmic linker associated protein 2 | 1.8 | 2E-29 |
| Chr14:106,498,501 | Intron | LINC01881 | long intergenic non-protein coding RNA1881 | 1.8 | 2E-29 |
| Chr6:129,416,001 | Intron | LAMA2 | laminin subunit alpha 2 | 1.8 | 4E-29 |
| Chr14:31,750,251 | Intron | NUBPL | nucleotide binding protein like | 1.8 | 2E-28 |
| Chr8:37,842,001 | Coding | ADGRA2 | adhesion G protein-coupled receptor A2 | -1.2 | 1E-12 |
| Chr1:44,724,501 | Coding | C1orf228 | chromosome 1 open reading frame 228 | -1.2 | 9E-13 |
| ChrX:46,575,001 | Coding | CHST7 | carbohydrate sulfotransferase 7 | -1.2 | 5E-13 |
| Chr14:104,729,501 | Coding | ADSSL1 | adenylosuccinate synthase like 1 | -1.3 | 1E-15 |
| Chr2:202,774,251 | Intron/promoter | ICA1L | islet cell autoantigen 1 like | -1.3 | 1E-15 |
| Chr1:155,190,001 | Coding | MUC1 | mucin 1, 2C cell surface associated | -1.4 | 2E-17 |
| Chr20:3,751,751 | Coding | HSPA12B | heat shock protein family A (Hsp70) member 12B | -1.4 | 8E-18 |
| Chr11:58,141,001 | Intron | OR9Q1 | olfactory receptor family 9 subfamily Q member 1 | -1.6 | 4E-22 |
| Chr1:45,803,751 | Coding | MAST2 | microtubule associated serine/threonine kinase 2 | -1.6 | 3E-23 |
| Chr7:636,001 | Intron | PRKAR1B | protein kinase cAMP-dependent type I regulatory subunit beta | -1.7 | 1E-24 |

$\#$ : High- versus average-risk value

Among the 69 differentially expressed genes, FAM83A and NEK2, showed the highest gene amplification rate (number of cases with gene amplification>200) and genetic alteration frequency (>10%), as well as overexpression in BC (*p*>0.001) and Oncoscore >50, as detected through METABRIC database, cBioportal, UALCAN and Oncoscore analyses FAM83A and NEK2 (**Table 1** and **Additional File 2: Fig. S2**) (17-19). The expression of FAM83A and NEK2 in the breasts of high- and average-risk women is shown in **Fig. 1D**. We detected a 4.5-fold increase in FAM83A and 2.2-fold increase in NEK2 expression in primary epithelial cells isolated from the breast of high-risk women when compared with cells isolated from breast tissue of average-risk women (**Fig. 1E**). Overexpression of both targets was detected also in a dataset of hTERT-immortalized epithelial cells as compared with the isogenic primary cells (20) (**Fig. 1F**). Moreover, immunostaining of a breast tissue microarray showed a 1.4 fold increase in FAM83A protein levels in the breast tissues from women at high risk of BC as compared with the breast tissues from subjects at average risk (*p*<0.0001, **Fig. 1G, Additional File 2: Fig. S3A** and **Additional File 1: Table S5**). FAM83A overexpression in normal breast tissues was associated with parity (*p*<0.001), tobacco use (*p*=0.01) and family history of BC (*p*=0.02) (**Additional File 1: Table S6**). On the contrary, NEK2 staining showed no difference in protein levels between the two groups (**Fig. 1G**). No difference in Ki67, estrogen receptor alpha (ERα), FOXA1 and GATA3 staining between high- and average-risk breasts was observed (**Additional File 2: Fig. S3B** and **Additional File 1: Table S5**). This data shows that FAM83A expression changes are specific to breasts of women at high risk of developing breast cancer.

### Genome-wide DNA methylation analysis reveals 1698 aberrant DNA methylation sites in normal breast tissue of high-risk women

With the goal of identifying alterations in regulatory regions leading to BC susceptibility, we performed a methylome analysis using the MethylCap-seq approach. Differential analysis of the methylated regions detected in the breasts from average-risk women and those from women at high risk of cancer revealed a wide chromosomal distribution of the epigenetic aberrations (**Fig. 2A**). DNA methylation changes with a ΔZ ≥1 (hypermethylated) or ≤-1 (hypomethylated) were selected. We identified 1698 regions methylated that differentiate the breast tissue of high-risk women from that of women at average risk (FDR□≤□5%), mapping to 1115 unique genes (**Additional File 1: Table S7**). The twenty most hypermethyated and hypomethylated regions are shown in **Fig. 2B** and **Table 2**. Interindividual variability in DNA methylation can be observed within each experimental group. 98.9% of the DNA methylation aberrations consisted of hypermethylated loci (*p*=□9□×□10^−□8^; **Fig. 2C**). More than 90% of hypermethylated loci localized in regulatory regions including the promoter, untranslated region, and introns, while only 41% of hypomethylated loci localized in these regions (**Fig. 2C**). However, hypomethylated regions were localized predominantly in the gene body (59%), a phenomenon that has been linked with the activation in cancers of aberrant intragenic promoters that are normally silenced (21, 22).

**Figure 2:**
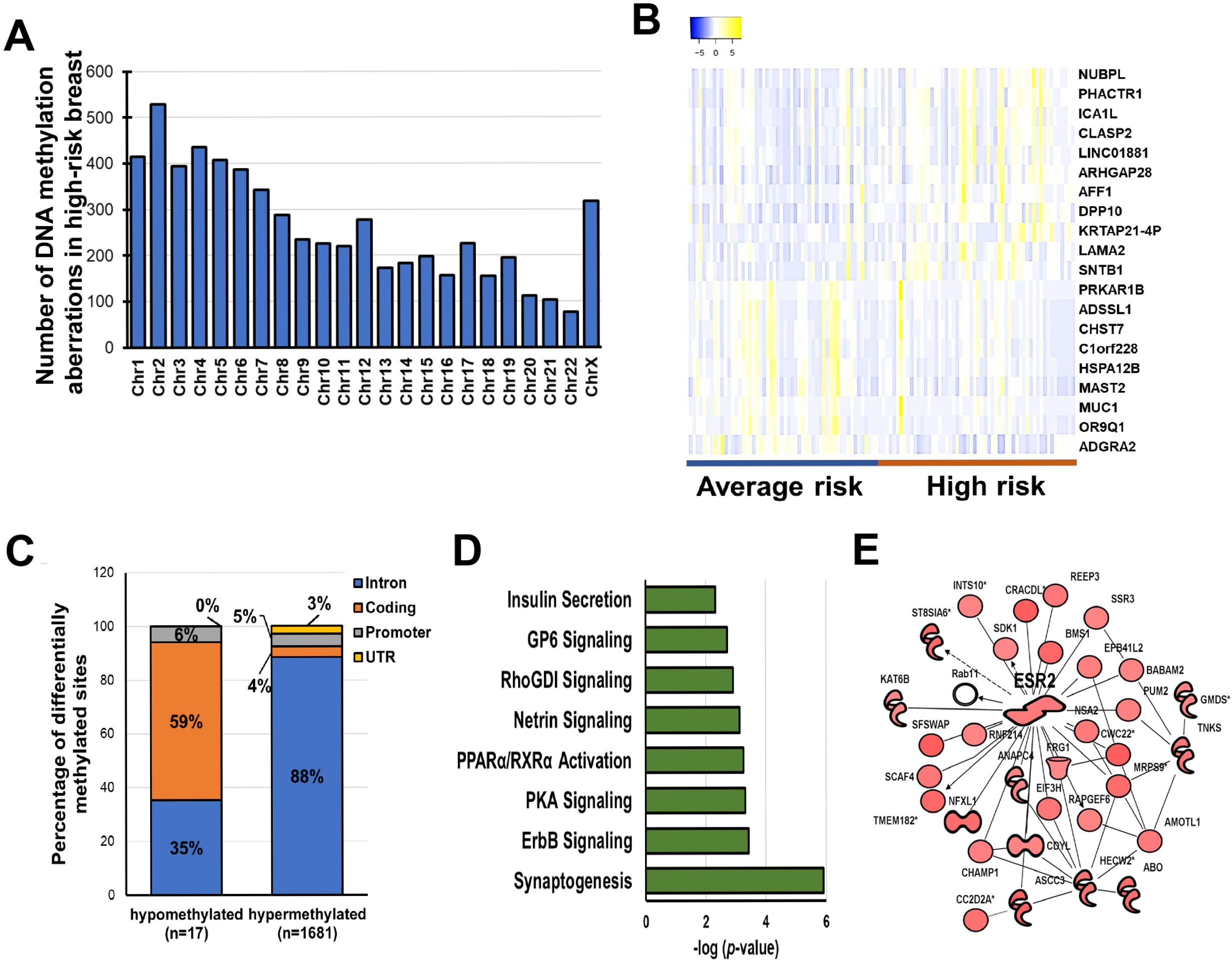
Methylome profiling of breast tissues from women at either high- or average risk of breast cancer. **A)** Chromosomal distribution of the DNA methylation aberrations observed in high-risk versus average-risk group. **B)** Heatmap of the 20 highest differentially methylated regions in high-risk breasts as compared with average-risk breasts at FDR<0.05. The overlapping gene name is indicated on the left. **C**) Genomic localization (intron, coding, promoter or UTR) of the DNA methylation aberrations including regions either hypo- or hyper-methylated in high-risk versus average-risk breasts. Data are shown as percentage of each genomic localizaton versus the total number of sites. **D)** Pathway analysis of the genes affected by DNA methylation aberrations (FDR<0.05) in high-risk breasts as compared with breast from women at average risk for breast cancer. **E)** One of the molecular networks including the genes affected by DNA hypermethylation.

Pathway analysis revealed the involvement of cell adhesion (aka synaptogenesis, *p*=1.2E-06), ErbB (*p*=3.7E-04) and protein kinase A (*p*=4.8E-04) signaling pathways (**Fig. 2D, Additional File 1: Table S8**). Notably, one of the molecular networks showed ESR2 as the central molecule (**Fig. 2E**). Although ESR2 expression decreased in high-risk breasts (fold change=0.82), the intronic ESR2 hypermethylation showed no inverse correlation with ESR2 expression (*r*=-0.03, *p*=0.4; **Additional File 2: Fig. S4A-C**). One of the hypomethylated genes, MUC1 (ΔZ=1.4, FDR=1.6E-17) is reported to be aberrantly overexpressed in over 90% of breast tumors (23, 24) (**Additional File 2: Fig. S4D**). However, no significant difference in MUC1 expression was observed between high- and average-risk breasts (**Additional File 2: Fig. S4E**). In the analyzed cohort, DNA methylation was not highly affected by either racial background or menopausal status of the tissue donors (FDR>0.05; **Additional File 1: Table S9**). Finally, we found overlap between DNA methylation aberrations in high-risk breasts and breast cancer-related DNA methylation signatures such as those identified by Saghafinia *et al* (4%, 25/666, (25)), Chen *et al* (6%, 10/174, (26)), de Almeida *et al* (9%, 25/285, (27)), and Xu et al (9%, 37/414, (28)) (**Additional File 1: Table S10**).

### DNA methylation and gene expression changes in high-risk breast show a weak correlation

To identify potential epigenetically regulated genes linked with BC risk, we performed a Pearson’s correlation test on paired DNA methylation and gene expression data (**Fig. 3**). Among the 69 genes in **Table 2**), the expression of eight genes was associated with aberrant intronic DNA methylation, including six genes showing a direct correlation (APELA, DIO2, FEZF2, LPAR3, UNC5D and PRSS51) and two genes (PROK2, and SULT1C2) with a negative correlation (**Fig. 3A**). Furthermore, among the DNA methylation aberrations in **Table 2**, only the intronic hypermethylation of PHACTR1 (ΔZ =1.88, FDR= 1.0E-31) was negatively correlated with PHACTR1 expression (fold change=0.77, FDR=0.006, *r*=-0.21) (**Fig. 3B**). However, the correlations identified were weak (*r*: −0.2,-0.5) suggesting that other regulator events (chromatin modifications, gene amplification, nucleotide variants), rather than DNA methylation aberrations, may be the determinants of the transcriptomic changes observed in the high-risk breasts as compared with average-risk breasts.

**Figure 3:**
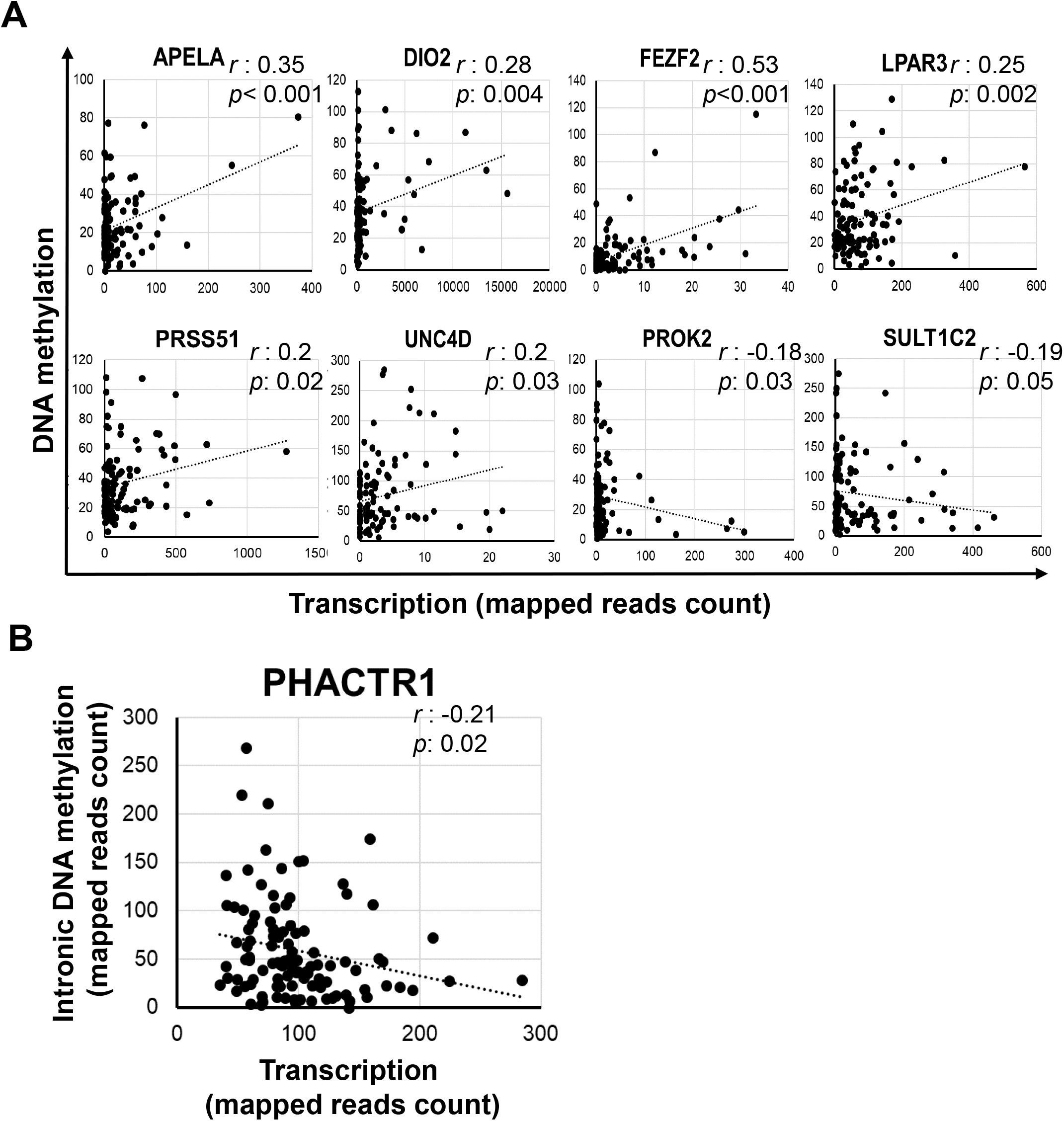
Correlation between degree of DNA methylation and gene expression. **A**) Pearson’s correlation analysis between DNA methylation value and expression of the genes found differentially expressed between high- and average-risk breasts. **B**) Pearson’s correlation analysis of the DNA methylation and expression of PHACTR1, hypermethylated in the breasts of high-risk women. *r* is the correlation coefficient and *p* is *p*value.

### Age-related molecular changes in cancer-free breast tissues in relation with cancer risk

Age is the strongest demographic risk factor for most human malignancies, including breast cancer. Age-related transcriptome and DNA methylation aberrations were investigated on breast tissues cores donated by 35 women at two separate time points (**Additional File 1: Table S11)**. Differential expression analysis (FDR<0.05) between the two donation time points revealed the dysregulation of 317 genes involved in LXR/RXR activation (*p*=7E-04), immune response (*p*=2E-03) and senescence (*p*=7E-03) (**Additional File 1: Tables S12 and S13**). Forty-eight age-related transcriptomic changes with a fold change (fc)≥2 and FDR <0.05 included two upregulated genes, CETP (fc:2.4;FDR=0.04) and HP (fc=2.3, FDR=0.03); and downregulation of five genes, SLC5A1 (fc=0.4; FDR=0.03), SLCO1A2 (fc=0.4; FDR=0.03), GRIA4 (fc=0.4; FDR=0.01), IL22RA2 (fc=0.4; FDR= 0.01), and CHRM1 (fc=0.4; FDR=0.03) (**Additional File 2: Fig. S5**). Furthermore, age-dependent dysregulation of the following five genes was enhanced in breast tissues from high-risk women: NEURL1, USP50, GRIA4, SPDEF and DNM3 (**Fig. 4A**). Notably, the expression of GRIA4 (*r*=-0.43, *p*=0.04) and DNM3 (*r*=-0.47, *p*=0.03) showed a negative correlation with their DNA methylation pattern, thus suggesting a potential epigenetic regulation for these two molecules (**Fig. 4B**).

**Figure 4:**
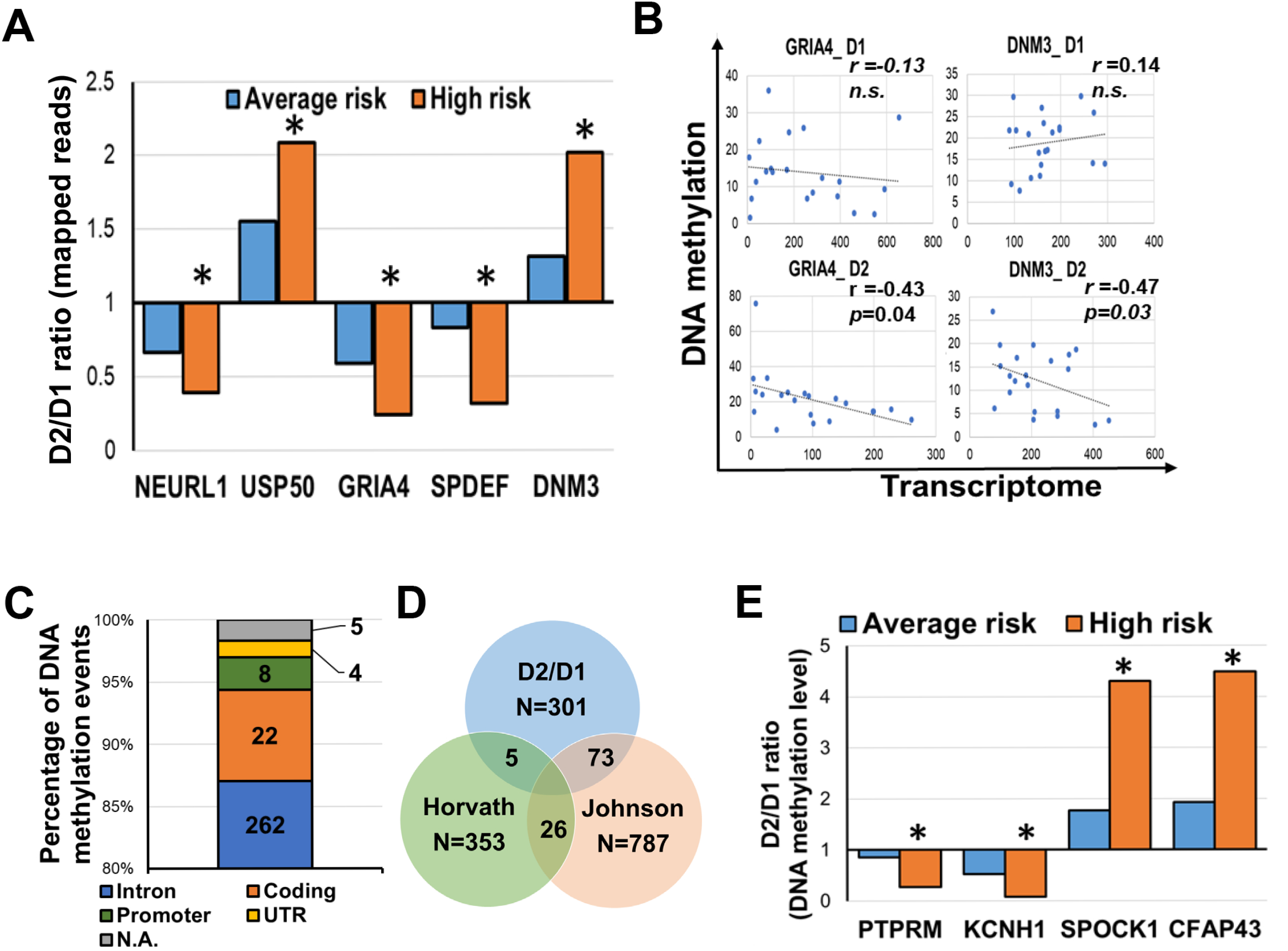
Age-related transcriptome and DNA methylation changes in healthy breast tissues. **A**) Differentially expressed genes between the first (D1) and second (D2) donation time point in the breast tissues from average (blue bars) and high- (orange bars) risk women. Ratio between D2 and D1 is shown. **B**) Pearson’s correlation test between DNA methylation and transcription of GRIA4 and DNM3 in average- and high-risk breasts at the two time points, D1 and D2, **C**) Number of genomic locations (intron, coding regions, promoter, UTR) of the age-related DNA methylation events. N.A.: not available. **D**) Venn diagram of the DNA methylation changes associated with age comparing our data set (D2/D1) with Horvath’ epigenetic clock (353 CpGs) or Johnson’s age-associated loci (787 CpGs) **E**) Differentially methylated regions between the first (D1) and second (D2) donation time point in the breast tissues from average (blue bars) and high- (orange bars) risk women. Ratio between D2 and D1 is shown. \**p*<0.05; \*\**p*<0.001

Age-dependent DNA methylation aberrations affected 301 loci corresponding to 280 unique transcripts (**Additional File 1: Table S14**). As previously reported (29), age-related DNA methylation alterations were predominantly hypermethylation events (85.4%) and affected the intronic regions (**Fig. 4C**). DNA methylation measurements were previously used to develop epigenetic biomarkers of aging, otherwise known as “DNA methylation age” or the “epigenetic clock” (30, 31). We observed a limited overlap between the 301 DNA methylation aberrations and the epigenetic clocks described by Horvath *et al* (1.4%, (31)), while 73 genes associated with the bin in our dataset overlapped with age-associated DNA methylation alterations reported by Johnson *et al* (24.2%,(29)) (**Fig. 4D**). Finally, we identified age-related DNA methylation aberrations enhanced in high-risk breasts, localized on four genes: PTPRM, SPOCK1, KCNH1, and CFAP43. (*p*<0.001, **Fig. 4E and Additional File 1: Table S14**). In contrast, both transcriptomic and DNA methylation aberrations affecting FAM83A and NEK2 resulted age-independent.

## DISCUSSION

This study aimed to define the distinct features of cancer-free breast tissues from women at high risk for BC and, thus, identify molecular markers that could potentially screen for women susceptible to cancer. We conducted transcriptome and methylome analyses using breast tissue cores donated by healthy women. The participants were divided into two cohorts based on their risk of developing breast cancer, according to the Tyrer-Cuzick lifetime risk assessment score: high-risk (≥20%) and average-risk (<20%) (32). Among the genes upregulated in high-risk breast, we identified two promising markers of BC susceptibility, FAM83A and NEK2. Furthermore, when investigating DNA methylation aberrations in high-risk breasts, we detected 4-10% overlap with cancer-related signatures.

Our transciptomic analysis of high- and average-risk breasts revealed significant changes in 69 genes (FDR<0.05). Pathway analysis suggested the activation of cell cycle and cell adhesion in the high-risk breasts. Furthermore, one of the molecular networks including the differentially expressed genes showed the involvement of FOXM1 signaling. FOXM1 itself is upregulated 1.6 fold in high-risk breasts (*p*=0.001).The transcription factor FOXM1 regulates the transcription of cell-cycle genes essential for exit from the G1/S phase into the G2/M phase such as cyclin A2, JNK1, ATF2 and CDC25A phosphatase as well as genes critical for chromosome segregation and cytokinesis (33). FOXM1 is overexpressed and plays critical role in tumorigenesis, metastasis, and drug resistance in a broad range of human cancer types, such as lung, gastric, and breast cancers (16). Compounds targeting FOXM1 expression or activity are under investigation (16). Our results suggest that the transcriptional dysregulation detected in high-risk breasts may be driven by FOXM1.

Two genes, FAM83A and NEK2, both upregulated in high-risk breast, showed a high Oncoscore (75.5 and 61.4, respectively), and have been reported amplified in breast cancer. FAM83A is the smallest member of the eight-member FAM83 family of proteins that share a conserved amino-terminal Domain of Unknown Function (DUF1669 domain) (34). It was identified based on its transforming potential (35-37). FAM83A upregulation has been detected in multiple human tumor types, including breast, lung, pancreatic and cervical cancer (37-44). Lee *et al* (45, 46) revealed the ability of FAM83A to confer resistance to epidermal growth factor receptor/ tyrosine kinase inhibitors (EGFR-TKIs) through interactions with c-RAF and PI3K p85 in breast cancer. The authors also showed that BC patients with high FAM83A expression had a worse prognosis. FAM83A depletion suppressed proliferation and invasiveness *in vitro* as well as tumor growth *in vivo* (36). Based on the aforementioned studies, FAM83A is considered a candidate oncogene and our findings suggest that FAM83A may be one of the first molecules dysregulated in cancer transformation. Our team is currently investigating the role of FAM83A in breast carcinogenesis. Moreover, our DNA methylation data, in agreement with previous literature, suggest that FAM83A overexpression is mainly driven by genomic amplification rather than epigenetic regulation (47, 48). Additional studies such as deep whole genome sequencing of DNA from breast tissues of high-risk women are required to support this hypothesis.

The NIMA-related kinase 2 (NEK2) protein belongs to a family of serine/threonine kinases, which have a role in mitotic progression by initiating the separation of centrosomes (49). NEK2 overexpression was previously reported in BC as result of gene amplification (47, 50). NEK2 depletion blocks cell cycle progression and tumor cell growth, making it an ideal therapeutic target (51). Notably, FOXM1 is reported to both bind NEK2 promoter and interact with NEK2 directly (52, 53). Our study further suggests a role of NEK2 dysregulation in breast carcinogenesis. However, we did not observe changes in NEK2 protein levels in breast tissues of high-risk women suggesting a disconnect between mRNA and protein levels, which is not uncommon. Further investigation of the role of NEK2 in breast carcinogenesis is needed.

We observed DNA methylation changes in high-risk breasts, consisting mostly of hypermethylation (98.8%) in the intronic regions (88%). Previous studies reported aberrant hypermethylation in normal breast tissue adjacent to the tumor (54). Hypermethylation in specific gene promoters is indeed linked to carcinogenesis through transcriptional silencing of tumor suppressor genes or regulatory regions within the genome leading to dysregulation of cell growth, cancer initiation and progression (55-57). We identified a 4-10% overlap between methylome aberrations in high-risk breasts and previously reported cancer-related signatures (25-28). The limited overlap may be linked to the different technical approaches (Methyl-capture vs Infinium HumanMethylation450 array) but may also suggest that cancer–related epigenetic marks are newly acquired during cancer initiation rather than being imprinted into the genome. Although the expression epigenetic modifiers such as DNMTs remain unaffected, we detected the upregulation in high-risk breasts of HASPIN (fc=1.7; FDR<0.005), a serine/threonine kinase involved into the phosphorylation of the histone H3 during mitosis (58), suggesting that other genetic and epigenetic mechanisms rather than DNA methylation may drive the transcriptomic aberrations in high-risk breasts.

Age is the strongest demographic risk factor for most human malignancies, including BC (59). The limited size of our cohort (N=35) prevented us from subdividing the subjects by age at tissue donation. Nevertheless, we identified age-related transcriptomic aberrations enhanced in high-risk breasts including GRIA4 and DNM3, which resulted potentially epigenetically regulated. In terms of DNA methylation aberrations, we found a limited overlap between the age-related DNA methylation changes here identified and epigenetic clock from Hovarth *et al*. (31) (**Additional File 1: Table S14**). However, a 24.2% overlap of our dataset with age-related DNA methylation aberrations described by Johnson *el at* (29) was detected. The limited overlap is probably due to the different platform used for DNA methylation detection (Infinium Human Methylation 450 array vs Methyl-Cap-seq) and the type of analysis (epithelium-specific deconvolution vs whole tissue)(29, 31). Notably, we identified specific age-related DNA methylation changes, located on PTPRM, KCNH1, SPOCK1, CFAP43 gene, enhanced in the high-versus average-risk breasts.

This study harbors some limitations: the relatively small sample size prevented us from investigating in details cancer-related varables such as racial background. The selection of normal breast tissue cores with high content in epithelial compartment limited the number of available samples (**Additional File 2: Fig. S6**). Outcome data for the women at high risk for BC is not available at this time; however, this cohort is under an ongoing annual medical follow up. Because of the faster processing time and smaller cost, we performed whole tissue analysis instead of the more epithelium-specific laser microdissection or single-cell analysis. This limits the compartment specificity of the data but generates a more comprehensive view of the molecular features of the entire breast tissue core. Further deconvolution analysis may overcome this limitation (60, 61).

## CONCLUSIONS

The present study reveals transcriptomic and epigenetic aberrations linked with BC risk. Transcriptional targets potentially promoting cancer susceptibility were identified. The described investigation provides an avenue for deciphering the functional relevance of genes involved in BC development and a rich resource for further investigation.

## METHODS

### Study cohorts

Breast specimens were obtained from the Susan G. Komen Tissue Bank at the IU Simon Comprehensive Cancer Center (KTB) and donated voluntarily upon informed consent by healthy women. Subjects were recruited under a protocol approved by the Indiana University Institutional Review Board (IRB protocols number 1011003097 and 1607623663). Subject demographics and breast cancer (BC) risk factors were collected using a questionnaire administered by the KTB and summarized in **Additional File 1: Table S1, S5** and **S11**. Breast tissue cores are collected by using a needle biopsy as previously described (14). The study cohort consisted of two groups: 1) For the transcriptome and methylome analyses, 146 women (median age: 39 years) were selected based on the lack of clinical and histological breast abnormalities and high content in breast epithelial compartment (cellularity>40%). Germline mutation status of the subjects was obtained from KTB. Data were retrieved from the LifeOmic’s Precision Health Cloud platform (https://lifeomic.com/products/precision-health-cloud/). Nine established breast cancer–predisposition genes (BRCA1, BRCA2, PALB2, ATM, CHECK2, BARD1, RAD51C, RAD51D, CDH1) were evaluated for variants identified as “pathogenic” or “likely pathogenic” in the ClinVar database (https://preview.ncbi.nlm.nih.gov/clinvar/) (**Additional File 1: Table S1**) (2, 3).

Thirty-five of these 146 women, including 10 at high risk and 25 at average risk for BC, donated their breast tissue at two time points at intervals from 1-10 years (mean: 3.2) between the tissue donations (**Fig. 1A** and **Additional File 1: Table S11**). 2) In a second analysis, paraffin-embedded breast tissue blocks related to 395 healthy women were obtained from the KTB and used to generate tissue microarrays. The cohort included 287 Caucasian, 66 African American, 49 Asian, with age ranging from 18 to 61 (**Additional File 1: Table S5**).

### Breast cancer risk assessment

Lifetime risk of developing BC was estimated by using the Tyrer-Cuzick risk score (IBISv8) (32) and a threshold of 20% to separate high- (≥20%) from average-risk (<20%) individuals. The Tyrer-Cuzick model was selected over the other risk estimation tools for its accuracy and inclusion of young subjects (62).

### Tissue processing and nucleic acid extraction

To limit stromal contamination, only breast tissue samples abundant in epithelial compartment (cellularity >40%) were selected and processed. Total DNA and RNA were isolated from fresh frozen breast tissue biopsies (80-1500mg) using AllPrep DNA/RNA/miRNA kit (Qiagen). Tissues were homogenized by using 2ml prefilled tubes containing 3mm zirconium beads (Benchmark Scientific, cat.# D1032-30), 350µl Lysis Buffer and 2-Mercaptoethanol, and BeadBug 6 homogenizer (Benchmark Scientific) in a cold room at the following conditions: 4000 rpm for 45 seconds was repeated twice with 90 seconds rest time. The concentration and quality of total RNA and DNA samples were first assessed using Agilent 2100 Bioanalyzer. A RIN (RNA Integrity Number) and DIN (DNA integrity number) of six or higher is required to pass the quality control.

### Whole transcriptome analysis

cDNA library was prepared using the TruSeq Stranded Total RNA Kit (Illumina) and sequenced using Illumina HiSeq4000. Data included 146 paired-end fastq sequence libraries (raw read length: 38 × 2). Reads were adapter trimmed and quality filtered using Trimmomatic ver. 0.38 (http://www.usadellab.org/cms/?page=trimmomatic) setting the cutoff threshold for average base quality score at 20 over a window of 3 bases. Reads shorter than 20 bases post-trimming were excluded. About 94% of the reads have both the mates passing the quality filters. Cleaned reads mapped to Human genome reference sequence GRCh38.p12 with gencode v.28 annotation, using STAR version STAR_2.5.2b (63). Only samples with about 99% of the cleaned reads aligned to the genome reference. Differential expression analysis was performed using DESeq2 ver. 1.12.3 (https://bioconductor.org/packages/release/bioc/html/DESeq2.html). Counts table containing mapped read counts for each gene, to be used as input for DESeq2 was generated using featureCounts tool of subread package (https://doi.org/10.1093/bioinformatics/btt656). Alternatively, we ran *t*-tests comparing the normalized read counts for the set of replicates from High risk samples to those for the set of replicates from Average risk samples. The normalized read counts were obtained from the DESeq2 run described above. The *p*values from the *t*-test were corrected for multiple testing using Benjamini-Hochberg method.

### DNA methylation analysis

Library was generated by using MethylCap Library Kit (Diagenode, Denville NJ, US) according to the manufacturer′s protocols followed by single-end 75-bp sequencing on Illumina Nextseq4000. Internal controls and duplicate samples were used to account for any batch effect and technical artifact. The data comprises of 146 paired end read libraries in FASTQ format. These libraries represent replicates for two samples - High risk (68 libraries) and Average risk (78 libraries). The libraries were sequenced across multiple runs and the combined read counts for each library were generated. Reads were adapter trimmed and quality filtered using Trimmomatic 0.38 (http://www.usadellab.org/cms/?page=trimmomatic) with the cutoff threshold for average base quality score set at 20 over a window of 3 bases. Reads shorter than 20 bases post-trimming were excluded. Approximately, 96% of the sequenced reads passed the quality filters to be considered “cleaned” reads. This quality control reduced the number of samples to 57 high- and 55 average-risk. Cleaned reads were mapped to Human genome reference GRCh38.p12 using BWA ver. 0.7.15 (64). Insert sequences were imputed from the concordantly mapped read pair alignments. More than 95% of the cleaned read pairs were concordantly mapped. A previously described differential methylation analysis using either Zratio or ΔZ (65, 66) was applied to the current methyl-capture dataset with a slight improvisation on the validation of the significance of differential methylation. For any given local bin of a given width on the genome, the method compares across samples, variation in deduplicated insert coverage distribution quantified as the bin’s z-score with respect to a larger genome region containing the bin. For this analysis, we used local non-overlapping bins with a fixed width of 250 bp with their z-scores computed relative to 25KB regions. Z-score is the number of standard deviations by which the bin coverage varies from the larger region’s mean coverage. A significant difference in Zscores, calculated as either as ΔZ or Zratio between the samples would indicate potential differential methylation for that bin, as previously described (67),. The analysis identified 159,438 bins, each 250bp wide, to be potentially differentially methylated between High risk and Average risk samples with z-ratios or ΔZ significant at 5% FDR and *p*-values from t-test ≤ 0.05. Based on positional overlap, these bins were annotated using annotation from gencode v28.

### Data analysis

Ingenuity Pathways Analysis (IPA, Qiagen, Redwood City, CA) was used for canonical pathway and molecular network analyses (68). Publicly available transcriptomic data from primary and immortalized breast epithelial cells were obtained from GEO (https://www.ncbi.nlm.nih.gov/geo/query/acc.cgi?acc=GSE108541) (20). Analysis of The Cancer Genome Atlas (TCGA) was performed by interrogating both cBioPortal (https://www.cbioportal.org/) and UALCAN (http://ualcan.path.uab.edu/) databases (69). Copy number variations (CNV) analysis was obtained from the interrogation of the Molecular Taxonomy of Breast Cancer International Consortium, METABRIC (17, 18). Oncoscore was used to rank genes according to their association with cancer, based on the available scientific literature (http://www.galseq.com/next-generation-sequencing/oncoscore-software/; accessed on 3/31/2021) (19).

### Primary breast epithelial cells and immunofluorescence

Primary breast epithelial cells were generated from cryopreserved breast tissue cores obtained from the KTB as previously described (14, 20). Immunofluorescence staining was performed as previously described (14). Briefly, 5,000 cells were cultures overnight into each well of an 8 well-chamber slide (BD Biosciences, San Jose, CA) and fixed with acetone: methanol (1:1) at −20°C for 10 min. After washing and blocking (PBS1X, 5% normal goat serum, 0.1%TritonX-100) steps cells were incubated with primary either rabbit anti-vimentin (Cell Signaling, D21H3, 1:100) or mouse anti-E-cadherin (Cell Signaling, 14472, 1:50) overnight. Upon three washes with PBS, cells were incubated with secondary antibodies (goat anti-mouse Alexa Fluor 568 or goat anti-rabbit Alexa Fluor 488; Thermo Fisher Scientific, 1:500) for 1 h at room temperature. After three washes with PBS, the coverslide was mounted using DAKO fluorescent mounting medium (S3023 Agilent, Santa Clara, CA) and the staining was visualized using a fluorescent microscope (Eclipse TS100, Nikon Instruments inc, Melville, NY).

### Quantitative real time polymerase chain reaction (qPCR)

Total RNA was extracted from cells using AllPrep DNA/RNA/miRNA kit (Qiagen). Reverse transcription was performed using SuperScript™ IV VILO™ Master Mix (Invitrogen cat#: 11756050) according to the manufacturer’s instructions. qPCR was performed using the TaqMan™ Universal PCR Master Mix (Applied Biosystems, cat# 4304437) and the following TaqMan Gene Expression Assays (Applied Biosystems/Thermo Fisher Scientific, Grand Island, NY): ACTB (Hs99999903_m1), FAM83A (Hs04994801_m1), and NEK2 (Hs00601227_m1). qPCR reactions were run on a StepOne Plus Real-Time PCR System (Applied Biosystems/Thermo Fisher Scientific) and data analyzed using the StepOne Software v2.3 (Applied Biosystems). Relative quantification was calculated with reference to ACTB and analyzed using the comparative C_T_ method. qPCR experiments were performed in triplicate.

### Tissue microarray (TMA) immunohistochemistry (IHC) analysis

Normal breast tissues microarrays from 683 women were generated from paraffin-embedded blocks obtained from the KTB at the Tissue procurement & Distribution core of the IU Simon Comprehensive Cancer Center. Due to loss of material during TMA construction and processing, 58% (n=395) of these tissue biopsies were interpretable. TMA was analyzed with the following antibodies FAM83A (Protein Tech 20618-1-AP, 1:100), NEK2 (MyBioSource MBS9607934, 1:100), Ki67 (DAKO IR 626, ready-to-use), estrogen receptor alpha (ERα) (clone:EP1, DAKO IR 084, ready-to-use), FOXA1 (Santa Cruz Biotechnology sc-6553, 1:100), and GATA3 (Santa Cruz Biotechnology sc-268, 1:50) (70). IHC was performed in a Clinical Laboratory Improvement Amendments (CLIA)-certified histopathology laboratory and evaluated by 3 pathologists in a blinded manner. Quantitative measurements generating positivity and H-score were done using the automated Aperio Imaging system using an FDA-approved algorithm (71).

### Statistical analysis

Comparisons between groups were done using either Student’s *t*-test or nonparametric Mann-Whitney test on GraphPad Prism 9. Difference between groups is considered significant at *p-*values<0.05. Pearson’s correlation analysis was performed to determine the strength and direction of the linear relationship between DNA methylation and transcription for given targets. Only correlations with a *p*<0.05 are shown. For transcriptome and methylome data, differential analysis was performed using DESeq2 and the previously described Z-score method (65, 66), respectively. *P*-values <0.05 are considered significanct and are corrected for multiple testing using the Benjamini-Hochberg False Discovery Rate (FDR) algorithm. For the tissue microarrays analysis nonparametric Wilcoxon rank-sum tests were used for unpaired analyses, as positivity and H-scores were not normally distributed, whereas nonparametric Wilcoxon signed-rank tests were used for paired analyses. The statistical software SAS version 9.4 (SAS Institute Inc., Cary, NC) was used to complete the statistical analyses with *p* < 0.05 considered significant. Baseline demographic characteristics were summarized as median (range) for continuous variables and number and percentage for categorical variables. Comparisons between groups were done using Chi-square tests (or Fisher’s Exact test, where appropriate) for categorical variables, or Wilcoxon test for continuous variables.

## LIST OF ABBREVIATIONS

BC: breast cancer
KTB: Susan G. Komen Tissue bank at IU Simon Comprehensive cancer center
IHC: immunohistochemistry
IPA: Ingenuity pathway analysis

## DECLARATIONS

### Ethics approval and consent to participate

Breast specimens were obtained from the Susan G. Komen Tissue Bank at the IU Simon Comprehensive Cancer Center (KTB) and donated upon informed consent by healthy women volunteers. Subjects were recruited under a protocol approved by the Indiana University Institutional Review Board (IRB protocols number 1011003097 and 1607623663)

### Consent for publication

Not applicable

### Availability of data and materials

Methylome and Transcriptome data are available in Gene Expression Omnibus (GEO) with GSE164694 (https://www.ncbi.nlm.nih.gov/geo/query/acc.cgi?acc=GSE164694) which includes the Sub Series GSE164640 (MeCap dataset) and GSE164641 (RNA-seq dataset). Transcriptomic data of primary and immortalized breast epithelial cells from Dr. Nakshatri’s team (20) were obtained from GEO with accession number GSE108541 (https://www.ncbi.nlm.nih.gov/geo/query/acc.cgi?acc=GSE108541).

### Authors’ contributions

N.M. conceived the idea and designed the experiments, analyzed, and interpreted the data, and was major contributor in writing the manuscript. N.M. and RG performed the experiments and generated the data. A.V. assisted with tissue processing. R.P, D.R., J.H., J.W., G.S., and S.A. analyzed the data. C.T. performed the immunostaining of the tissue microarray. P.R. provided the specimens used in this study. K.N. and H.N. provided input on the manuscript. A.M.S. participated in hypothesis generation, funded the work, and contributed to the manuscript preparation. All the authors read and approved the manuscript.

## Acknowledgements

Samples from the Susan G. Komen Tissue Bank at the IU Simon Comprehensive Cancer Center were used in this study. We thank the tissue donors, whose help and participation made this work possible. We thank the IU Simon Comprehensive Cancer Center for the use of the Tissue Procurement & Distribution Core, which provided the generation of the tissue microarray. We thank the Immunohistochemistry Core for the immunostaining and analysis of the tissue microarrays and the Center for Genomics and Bioinformatics Core at IU Bloomington for the –omics analyses.

## ADDITIONAL FILES

Additional File 1: Supplementary Tables. File format:.xls. It includes subjects demographics and raw data in form of tables.

Additional File 2: Supplementary Figues. File format:.pdf. It includes additional data related to the main findings shown in the main figures.

