## Supplementary Figures 1-6 for "Aberrant epigenetic and transcriptional events associated with breast cancer risk"

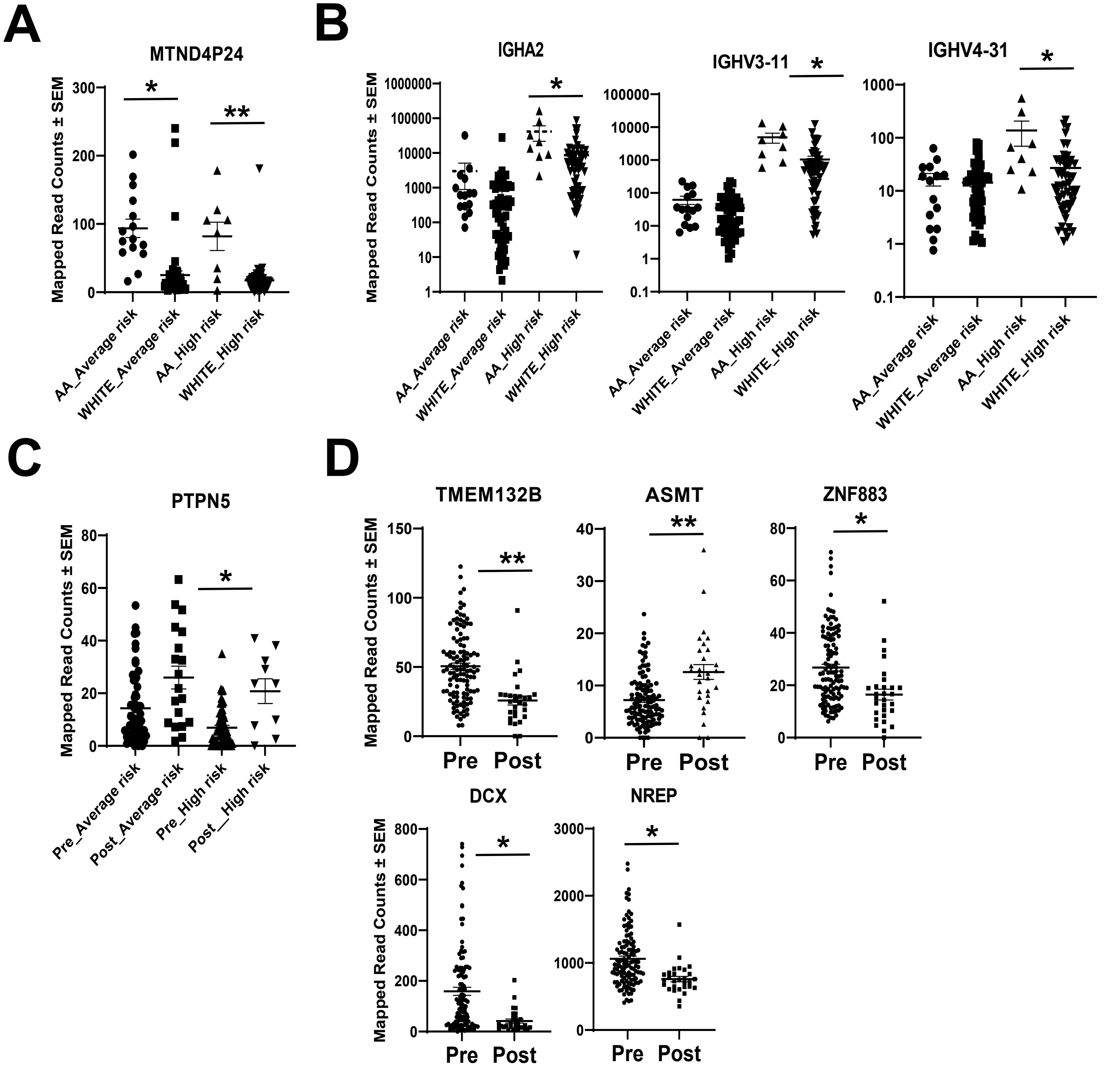


**Figure S1**: **Transcriptional changes in relation to racial background and menopausal status**. **A**) Pseudogene overexpressed in the breast from African American (AA) as compared with Caucasian (C) women (FDR=0.03). **B**) Transcripts differentially expressed between AA and C in high risk breasts only. **C**) Gene overexpressed in the breast from postmenopausal (Post) as compared with those from premenopausal (Pre) women in high risk women (FDR=0.02). **D**) Genes differentially expressed between Pre and Post samples independently from the risk score. Mapped read counts ± standard deviation (SEM) for each transcript are shown. Two-tailed unpaired t test was used to calculate the *p*value followed by Benjamini-Hochberg correction to obtain the FDR value. *:FDR<0.05; **:FDR<0.005

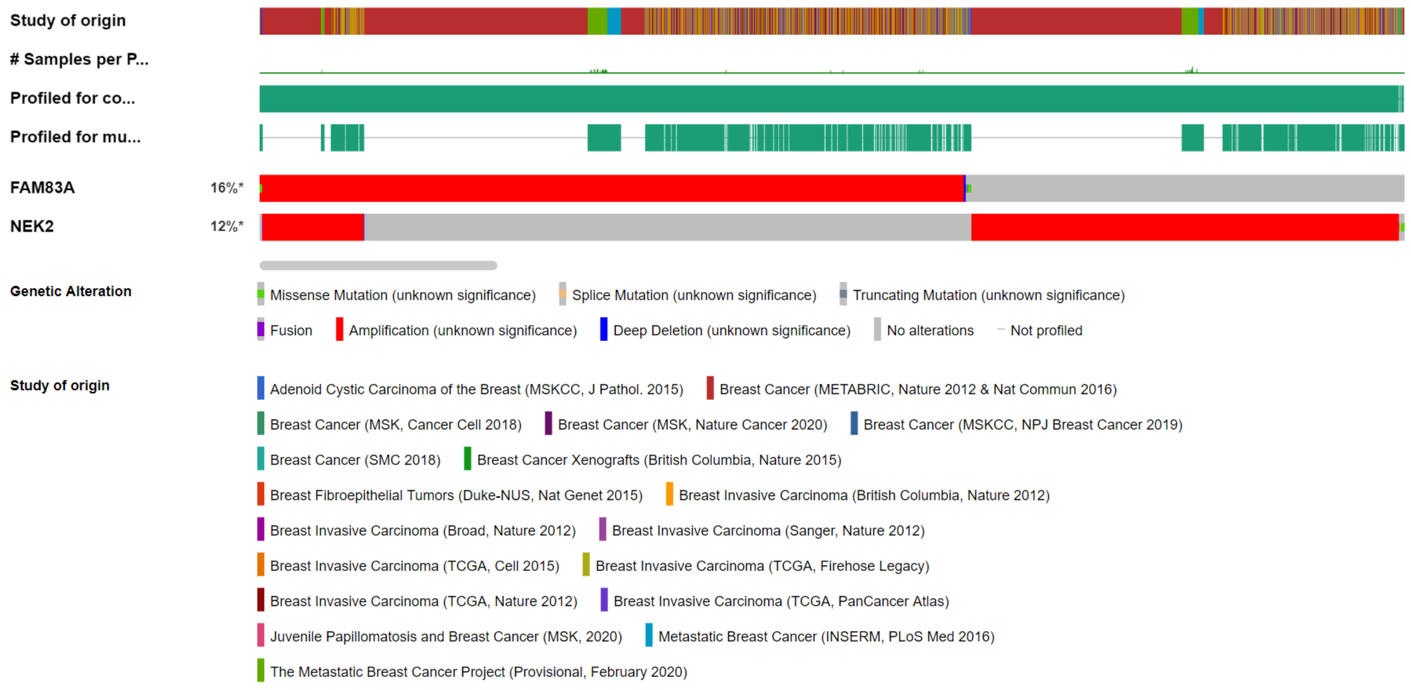


**Figure S2**: **FAM83A and NEK2 genetic alterations in breast cancer.** OncoPrint data for FAM83A and NEK2 were obtained by interrogating cBioPortal database.

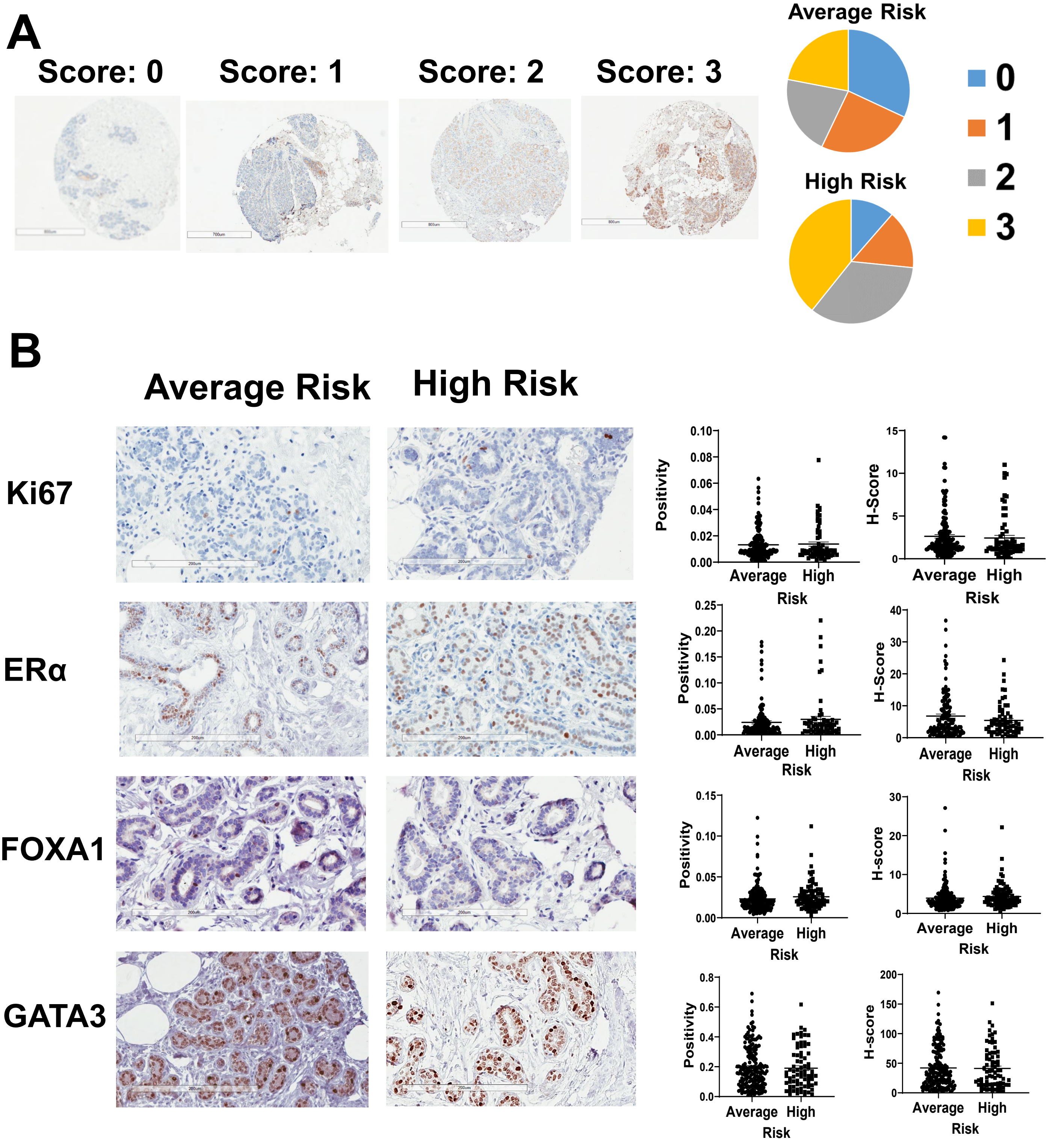


**Figure S3**: **Immunochemistry (IHC) stainings on breast tissue sections from women at either high or average risk for breast cancer..** A) Intensity of FAM83A IHC staining was evaluated (negative =0, borderline, minimal= 1, moderate=2, or severe=3. Left panel: representative images at 4x magnification; right panel: pie chart showing the distribution of the different staining intensity among the average risk (top) and high risk (bottom) breast tissue sections. **B**) representative images (left) and staining quantification for the indicated markers (right) in breasts from women at either high or average risk for breast cancer..

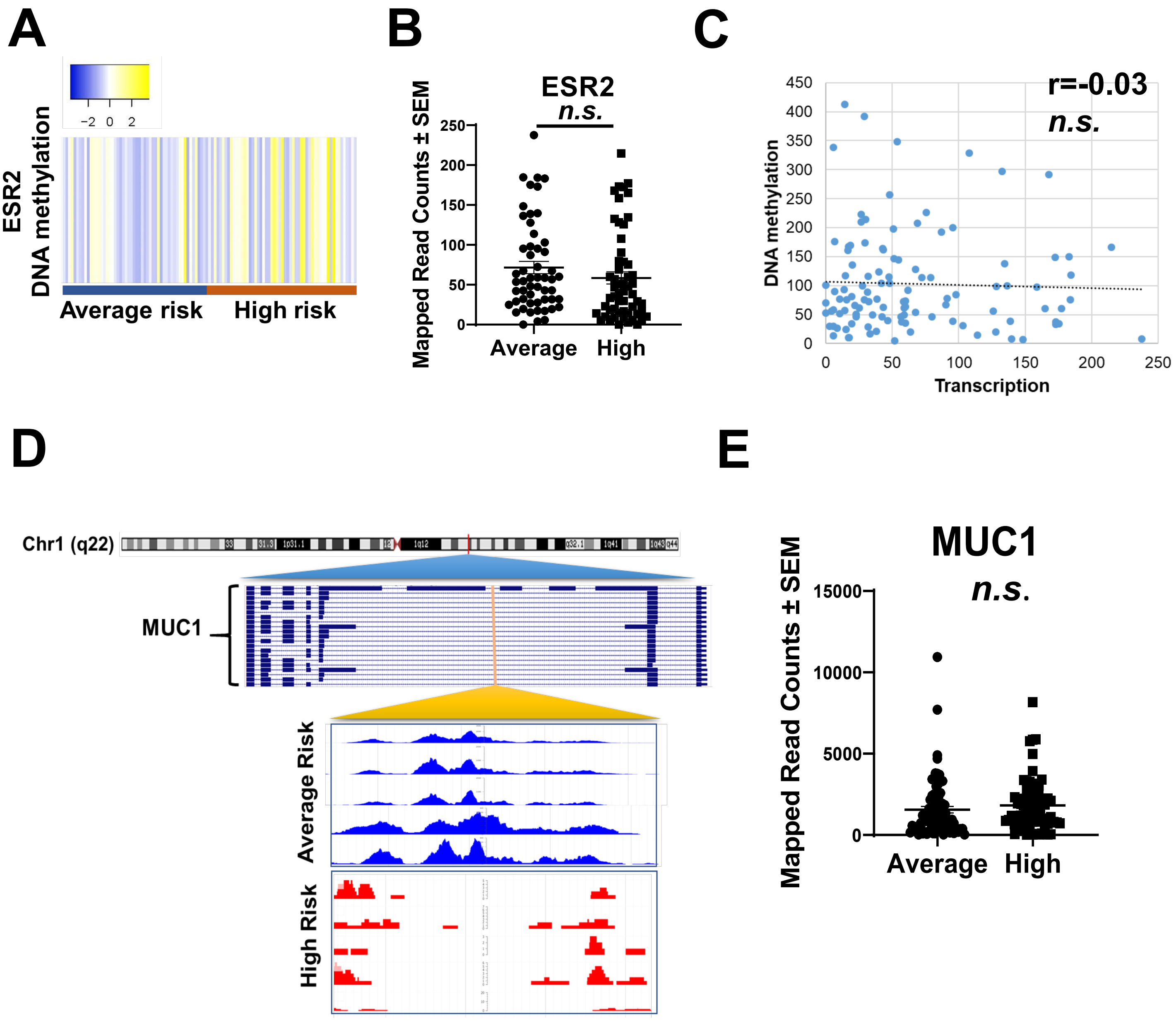


**Figure S4: ESR2 and MUC1 expression and methylation in breasts from women at either high- or average-risk for breast cancer**. **A**) Heatmap of the DNA intronic methylation status of ESR2 in average- and high-risk breasts. **B**) Expression of ESR2 in average- and high-risk breasts. Mann-Whitney unpaired non parametric test was used to determine the difference between the groups. **C**) Pearson correlation test between the DNA methylation and expression of ESR2. **D**) Genomic location of MUC1 gene and the CpG sites affected by DNA methylation aberration in high-risk breasts. The magnification shows representative images of the DNA methylation status of MUC1 intronic region in the breast of high-risk (red) and average-risk (blue) women. **E**) MUC1 expression in high- and average-risk breast tissues. Statistical significance is calculated with Whitney- Mann test using GraphPad Prism v9. *n.s.*: not significant

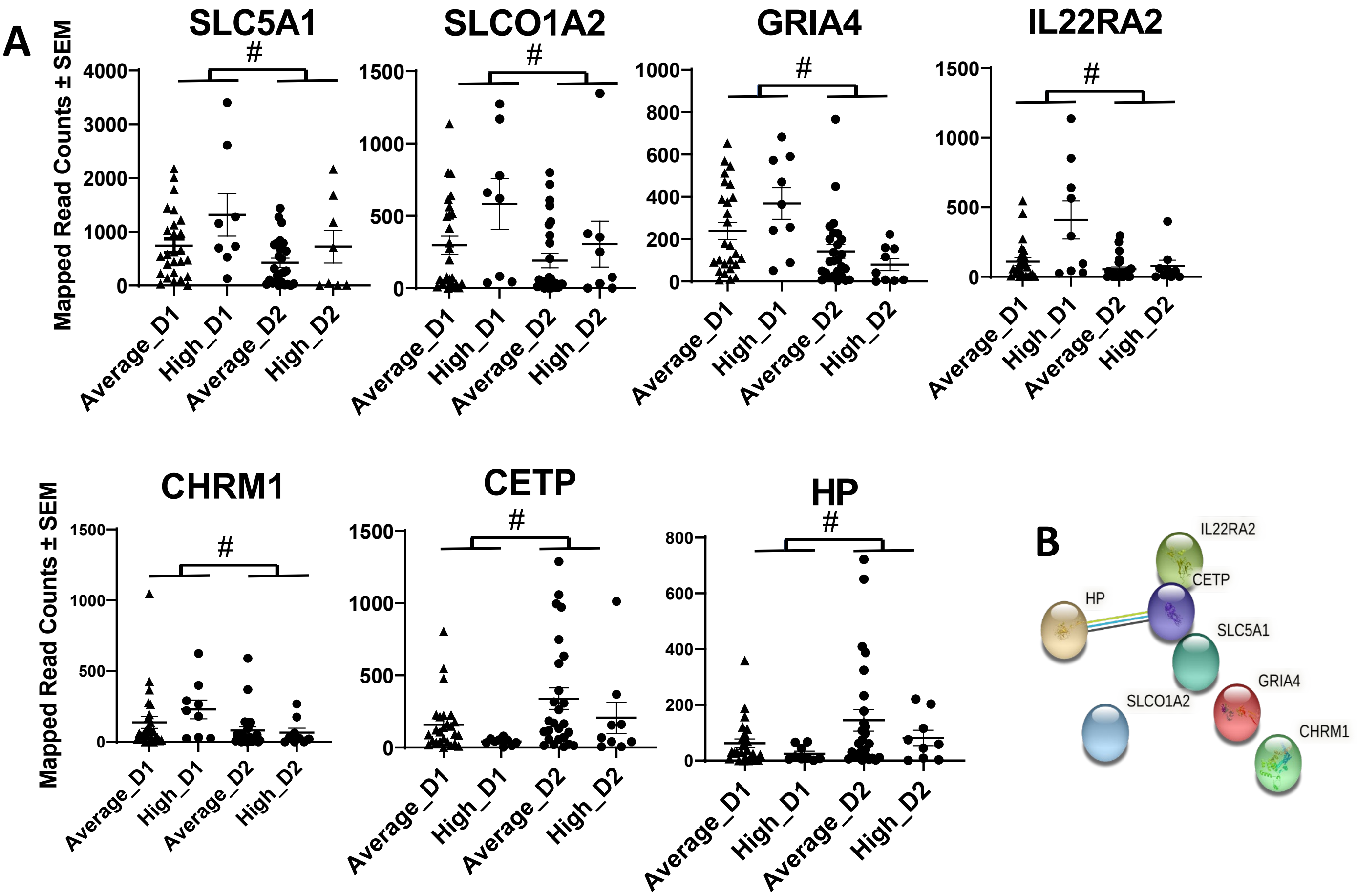


**Figure S5**: **Age-related transcriptome changes in healthy breast tissues**. **A**) Expression of the genes differentially expressed between the first (D1) and second (D2) donation time. Data are expressed as mean ± SEM. **B**) STRING inter-molecular interaction analysis between the genes differentially expressed between the two donations. # FDR<0.05

**A**

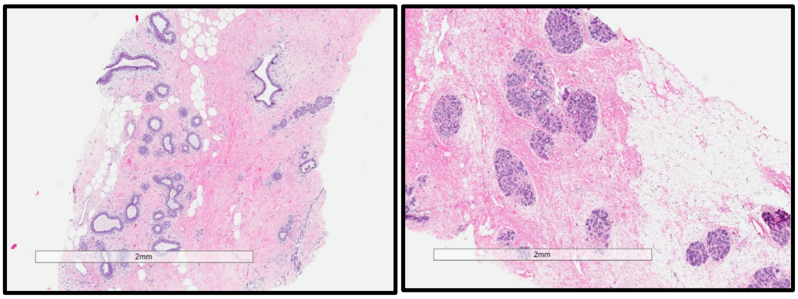


**B**

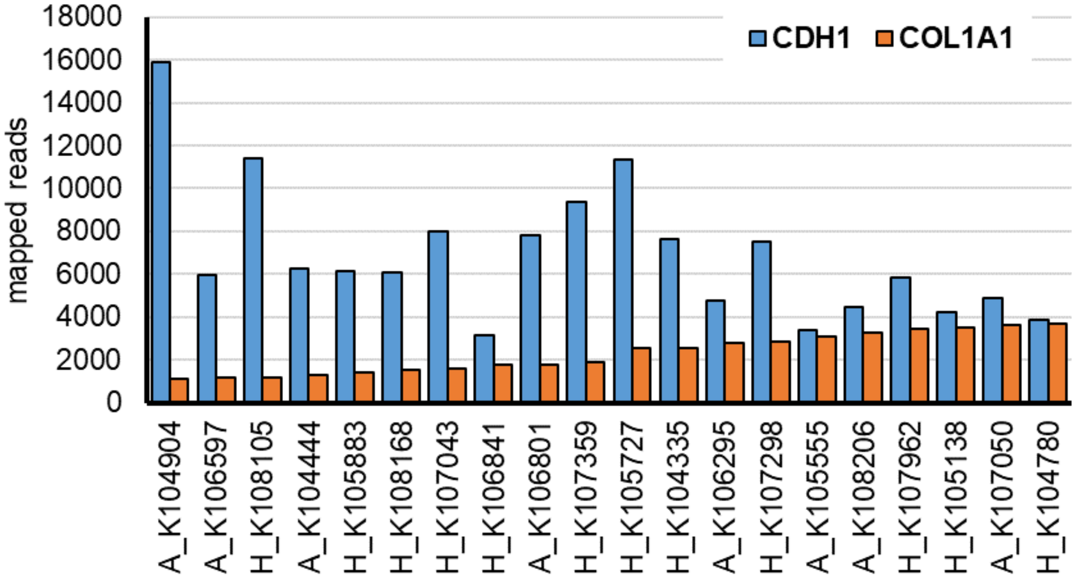


**Figure S6**: **Epithelial and stromal composition of the normal breast tissues**. **A**) Representative hematoxylin & eosin images of the normal breast tissues used in this study. **B**) Expression level of specific markers for epithelial cells (CDH1) and fibroblasts (COL1A1) in 20 representative breast samples shows the heterogeneity in the content of breast epithelium versus the stroma, although the first is more abundant in the majority of the samples.
